# Anthropometric and functional profiles of elite athletes from different categories at ABADÁ Capoeira World Championship

**DOI:** 10.1101/330126

**Authors:** Lucas G Almeida, Eduardo S Numata Filho, Geovani A dos Santos, José T C Cardoso - Mestre Camisa, Sérgio R Moreira

**Affiliations:** College of Physical Education, Federal University of Vale do São Francisco – Univasf, PE, Petrolina, Brazil.; Graduate Program on Physical Education, Federal University of Vale do São Francisco – Univasf, PE, Petrolina, Brazil.; Founder president of Associação Brasileira de Apoio e Desenvolvimento da Arte Capoeira – ABADÁ Capoeira. Rio de Janeiro, RJ, Brazil.

**Keywords:** body composition, motor performance, athletes, martial arts

## Abstract

This study aimed to characterize the anthropometric and functional profiles of 50 male elite competitors at the 2017 Capoeira World Championship organized by ABADÁ Capoeira School in the city of Rio de Janeiro. The competitors were divided into different weight categories: lightweight (VIOLA, ≤76.9 kg; n = 15), intermediate weight (MEDIO, 77.0–85.9 kg; n = 25), and heavyweight (GUNGA, ≥86.0 kg; n = 10). Two evaluation batteries were performed: 1) anthropometry and somatotype determination and 2) functional performance on push-up, sit-up, and sit-and-reach tests; quadrant jump test (QJT); squat jump (SJ); and countermovement jump (CMJ). Results reveal that the mesomorphic component of the somatotype differed between the GUNGA subgroup and other groups (F_[2,47]_ = 7.617; p = 0.001), while ectomorphy differed between the VIOLA and GUNGA subgroups (F_[2,47]_ = 3.899; p = 0.027). The “endo-mesomorph” classification predominated in the three investigated categories. For functional performance, there was a difference in QJT between the VIOLA and GUNGA subgroups (F_[2,47]_ = 4.299; p = 0.019). The endomorphism had a negative correlation (p < 0.01) with the performance in the sit-up (r = -0.51), push -up (r = -0.39), SJ (r = -0.45), and CMJ (r = -0.49). We concluded that at the international level, male elite competitors show predominance in the mesomorphic component of the somatotype. Furthermore, the endomorphic component correlated inversely with functional performance in trunk and upper limb resistance tests and lower limb explosive strength tests. These results can help coaches in targeting specific training programs for Capoeira athletes who aim for a high competitive level.

## Introduction

Capoeira is defined as an athletic performance that comprises an attack and defense system with individual character. It is considered a martial art and one of the most important popular manifestations in Brazil, being the only sport recognized by the *United Nations Educational, Scientific and Cultural Organization* (UNESCO) as Intangible Cultural Heritage of Humanity [1]. It is currently practiced in >150 countries across the five continents [2] and by different social groups [3]. This martial art has been influenced from several historical periods since the sixteenth century and includes sport, education, and artistic, musical, and folkloric aspects in its play [4].

Due to its popularity, Capoeira has been focused as an important study subject in the academic environment [3]. Monteiro et al. (2015) [5] have verified the reaction time among Capoeira practitioners with different experience times in the modality. Maia et al. (2010) [6] have compared anthropometric and cardiovascular variables among Capoeira practitioners and sedentary individuals. Additionally, some studies also have demonstrated that the practice of Capoeira can chronically reflect cardiovascular [7] and neuromuscular benefits [8] in beginner practitioners.

In contrast, the shape of the body and the distribution of the body composition have become increasingly distinct among sports but similar in each of them. This assumption suggests that each sport attributes an ideal physical characteristic to its practitioner. This anthropometric trait can be associated with functional performance, which also needs to be clarified in different sports [9]. The somatotype of athletes in different martial arts, such as jiujitsu [10], judo [11], taekwondo [12], boxing [13], karate [14], and Greco-Roman wrestling [15], is well documented in the literature. However, considering the possible requirements in the Capoeira modality, studies on the physical and functional profiles of its practitioners are still necessary.

Since a few decades, Capoeira has been managed by different associations, which are commonly denominated groups or schools [3]. From this perspective, there is the ABADÁ Capoeira School, which was founded in 1989 in Rio de Janeiro, and today, it has >40,000 members distributed in >60 countries worldwide [16]. The ABADÁ Capoeira School organizes biennially the Capoeira World Championship, which classifies the best athletes from an integral methodology evaluating batteries with different rhythms of the Capoeira game. Moreover, the athletes compete in categories differentiated by technical level and body weight. In this way, this study aimed to characterize the anthropometric and functional profiles of elite athletes from different categories and competitors in the world championship of the ABADÁ Capoeira School. This study hypothesized that anthropometric, but not functional, profile may differ between body weight categories. Such information can contribute to a better segmentation of specific training programs in Capoeira, besides allowing an understanding of the characteristics of the physical performance in a specific requirement in the modality.

## Materials and methods

### Subjects

The present study was conducted in accordance to the requirements stipulated in the Declaration of Helsinki and approved by the Research and Ethics Committee of the Federal University of Vale do São Francisco (protocol 2.210.868 CEDEP). After signing an informed consent form, 50 apparently healthy male athletes were examined for this investigation. All evaluations occurred during the 11th Capoeira World Championship of the ABADÁ Capoeira School in August 2017 in the state of Rio de Janeiro at the *Mestre Bimba Educational Center* (*CEMB*). The sample was subdivided into three weight categories defined by the competition rules. The light category (VIOLA, ≤ 76.9 kg; n = 15); intermediate category (MEDIO, 77.0–85.9 kg; n = 25), and heavy category (GUNGA, ≥ 86.0 kg; n=10). All the competitors that were evaluated belonged to category A (higher technical level), which composes the upper rope graduates in the competition (purple, purple-brown, brown, and brown-red). To meet the aim of the work, two evaluation batteries were set up: 1) anthropometry and somatotype determination and 2) execution of functional performance tests. Comparisons of the variables among the three different weight categories, as well as somatotype correlations with functional performance, were analyzed.

The purpose of the cardiovascular parameter evaluation was only to mark the different subgroups regarding their cardiac characteristics during the competition period. The blood pressure was verified by an automatic monitor (*Microlife®* model BP 3AC1-1 PC, Widnau, Switzerland). The heart rate and R-R intervals were recorded by the Polar H7 heart rate belt transmitter, Polar Electro Oy [17], connected via Bluetooth in a smartphone managed by the app *Elite HRV* [18]. All analyses were run through the Kubios HRV analysis software version 3.0 (Biosignal Laboratory, University of Kuopio, Kuopio, Finland).

### Anthropometric evaluation

The anthropometric evaluation followed the guidelines of the International Society for the Advancement of Kinanthropometry [19]. Body mass (EKS 9824 digital scale, 0.1-kg variation), height, and circumferences of the waist, hip, thigh, leg, relaxed arm, and flexed arm were measured by a flexible steel tape with sequential scale, resolution in millimeters, with 2 meters long and 6 mm wide (Cescorf, Porto Alegre/RS, Brazil). Body mass index (BMI) was calculated through the equation weight × height^2 (-1)^ and the waist-to-hip ratio (WHR) through the waist × hip^-1^ equation. The skinfolds of the triceps brachii, subscapular, supraspinal, abdominal, medial thigh, chest, and calf regions were evaluated in triplicate using a skinfold caliper with sensitivity of 0.1 mm, total amplitude of 85 mm, and pressure of 10 g × mm^2 (-1)^ (Cescorf/Mitutoyo, Porto Alegre/RS, Brazil). The biepicondylar humerus and femur breadths were measured using a stainless-steel bone caliper that was 16 cm in length with resolution in millimeters (Cescorf, Porto Alegre/RS, Brazil). The body fat (BF) percentage was obtained from Jackson and Pollock’s predictive equation [20] and corrected for the Cescorf skinfold caliper [21]. Finally, the somatotype was determined and represented in the somatochart (endomorphy, mesomorphy, and ectomorphy), following the guidelines of Heath and Carter [22].

### Functional evaluation

Immediately after the anthropometric measurements, several motor tests for the functional evaluation of the Capoeira athletes were set up. The performance tests were conducted in the following order: 1-min push-up test (push-up), 1-min sit-up test (sit-up), sit-and-reach test, squat vertical jump performance test (SJ), countermovement vertical jump performance test (CMJ), and quadrant jump test (QJT). The subjects were instructed to execute the best performance in each of the tests above. The sit-and-reach test, SJ, CMJ, and QJT were performed three times with a 15-s interval between each trial, where the best performance was considered in the evaluation.

The push-up evaluated the strength and resistance of the upper limbs muscles, where the volunteers were placed in the ventral decubitus positions, with their hands aligned to the pectoral region and 1– 2 cm away from the shoulders. The initial position consisted of the elbows extended, and a repetition was considered when the volunteer’s chest approached the ground and returned to the initial position. The objective was to perform the maximum number of repetitions in 1 min [23].

The sit-up evaluated the strength and resistance of the abdominal muscles, where the volunteers were in the dorsal decubitus position. Their feet were supported on the ground. The knee and hip articulations were 90° flexed with the arms crossed and hands resting on the shoulders. The subjects were instructed to perform a trunk flexion until the elbows touched the anterior portion of the thighs. The objective was to perform the maximum number of repetitions in 1 min [23].

The sit-and-reach test evaluated the range of motion of the joints of the posterior muscle chain (triceps sural, hamstrings, and gluteal and spinal erectors). The sit-and-reach box proposed by Wells (Sanny®, Curitiba/PR, Brazil) was used for this test. The dimensions of the box were 30.5 cm × 30.5 cm × 30.5 cm with 26 cm long scale in its extension. The zero point was at the closest extremity to the subject, and the 26-cm scale coincided with the place where the feet were supported. The volunteers were instructed to sit on the ground and position their feet on the indicated place of the box with the knees extended. The knees were completely extended during the execution, the hands were overlapped, and the elbows were extended. The objective was to flex the trunk to advance the hands as much as possible in the measurement scale [24].

The SJ and CMJ evaluated the explosive and elastic forces of the lower limb muscles by a specific jump plate (Cefise®, São Paulo/SP, Brazil) with interface to a microcomputer, where, through the Jump System Pro software version 1.0, it was possible to manage the measures. The subjects performed the jump after 3 s in a semi-squat position for the SJ. However, in the CMJ, the subjects executed the jump starting from the standing position to a semi-squat and using the stretch-shortening cycle. During both vertical jump tests, the volunteers’ hands were positioned on their waist region. The results obtained in the vertical jump tests made the calculation of the elasticity index (EI) from the following equation possible: EI = [(CMJ - SJ) / SJ * 100].

The QJT evaluated the agility from the displacement of the subjects over a cross-marked adhesive tape on the ground, which had a 1 m length by 1 m width. With the feet together, the volunteers jumped through the quadrants as quickly as possible in 10 s. The number of correct jumps executed was evaluated [25].

### Statistical Analysis

Data are expressed as mean, standard deviation, coefficient of variation, and minimum and maximum values. The test of normality, using the Shapiro-Wilk test, was conducted. One-way ANOVA was used to compare the study subgroups. Mauchly’s test was used to verify the sphericity of the data; in case of violation, the Greenhouse-Geisser correction was adopted. The Tukey post hoc test was applied to identify the pairs of differences. Pearson’s linear correlation coefficient was calculated to quantify the degree of association between anthropometric and functional variables. The level of significance was set at p-value < 0.05, and all statistical analyses were conducted in the SPSS 22.0 software.

## Results

Table 1 shows the general characteristics mentioned and results obtained in the anthropometric variables of the elite Capoeira athletes. Significant differences in body mass (F_[2.47]_ = 57.577; p < 0.001) and BMI (F_[2.47]_ = 17.650; p < 0.001) were found among the three subgroups. For the height, the VIOLA subgroup presented lower values compared to the MEDIO and GUNGA subgroups (F _[2,47]_ = 10.435; p < 0.001). The systolic blood pressure was significantly different (F_[2,47]_ = 4.662; p = 0.014) between the VIOLA and GUNGA subgroups.

**Table 1.** Anthropometric and cardiovascular characterization of the elite Capoeira athletes in the three categories (n = 50).

|  | VIOLA (n = 15) |  |  | MEDIO (n = 25) |  |  | GUNGA (n = 10) |  |  |
| --- | --- | --- | --- | --- | --- | --- | --- | --- | --- |
| | $\bar{X} \pm SD$ | Range | CV (%) | $\bar{X} \pm SD$ | Range | CV (%) | $\bar{X} \pm SD$ | Range | CV (%) |
| **Age (years)** | 32.8±7.2 | (19.0-44.0) | 30 | 34.3±5.3 | (24.0-49.0) | 16 | 36.1±4.8 | (30.0-46.0) | 13 |
| **PT (years)** | 20.7±6.2 | (11.0-35.0) | 9 | 21.3±4.6 | (15.0-35.0) | 22 | 24.3±6.3 | (18.0-33.0) | 21 |
| **BM (kg)** | 71.1±6.3* | (57.3-76.9) | 4 | 81.8±2.9† | (77.0-85.9) | 5 | 93.6±6.0 | (86.0-106.9) | 7 |
| **Height (cm)** | 168.0±7.1* | (155.0-184.0) | 8 | 174.0±4.5 | (167.0-182.0) | 3 | 178.0±4.4 | (170.0-184.0) | 2 |
| **BMI (kg/m<sup>2(-1)</sup>)** | 25.0±1.9* | (21.6-28.5) | 8 | 27.0±1.5† | (24.0-29.0) | 6 | 29.4±1.9 | (26.0-33.3) | 7 |
| **WHR** | 0.88±0.03 | (0.83-0.94) | 4 | 0.86±0.03 | (0.81-0.95) | 3 | 0.86±0.02 | (0.83-0.90) | 3 |
| **BF (%)** | 9.8±3.4 | (4.9-15.6) | 35 | 11.9±3.6 | (6.7-20.0) | 30 | 12.5±5.9 | (5.8-20.8) | 47 |
| **SBP (mmHg)** | 121.0±6.7† | (112.5-135.5) | 6 | 124.4±11.3 | (104.5-146.0) | 9 | 133.9±12.8 | (117.0-162.0) | 10 |
| **DBP (mmHg)** | 74.6±9.3 | (60.0-89.5) | 13 | 76.7±5.9 | (66.0-88.5) | 8 | 79.9±8.4 | (70.0-98.0) | 11 |
| **HR (bpm)** | 71.8±9.4 | (60.8-95.7) | 13 | 74.5±10.0 | (52.0-93.8) | 13 | 72.4±5.4 | (64.0-80.5) | 8 |
| **RRi (ms)** | 846.5±98.5 | (626.6-986.7) | 12 | 820.3±118.8 | (639.1-1152.3) | 14 | 832.6±64.1 | (745.2-938.0) | 8 |

**Note:** $\bar{X}$ = mean; SD = standard deviation; CV = coefficient of variation; PT = practice time; BM = body mass; BMI = body mass index; WHR = waist-hip ratio; BF = body fat; SBP = systolic blood pressure; DBP = diastolic blood pressure; HR = heart rate; RRi: R-R interval. \* $p < 0.05$ to *MEDIO* and *GUNGA*; † $p < 0.05$ to *GUNGA*

Considering the functional evaluation, significant differences (F_[2,47]_ = 4.299; p = 0.019) for QJT were observed between the VIOLA and GUNGA subgroups (Table 2).

**Table 2.** Functional characterization of the elite Capoeira athletes in the three categories (n = 50).

|  | VIOLA (n = 15) |  |  | MEDIO (n = 25) |  |  | GUNGA (n = 10) |  |  |
| --- | --- | --- | --- | --- | --- | --- | --- | --- | --- |
| | $\bar{X} \pm SD$ | Range | CV (%) | $\bar{X} \pm SD$ | Range | CV (%) | $\bar{X} \pm SD$ | Range | CV (%) |
| **Push-up (reps)** | 37.0±11.2 | (20.0-64.0) | 30 | 39.6±9.1 | (22.0-57.0) | 23 | 38.9±12.4 | (23.0-59.0) | 32 |
| **Sit-up (reps)** | 41.2±6.6 | (25.0-52.0) | 16 | 41.6±8.8 | (21.0-53.0) | 21 | 34.6±8.7 | (22.0-50.0) | 25 |
| **Sit-and-reach (cm)** | 39.8±4.8 | (33.5-49.0) | 12 | 40.4±5.0 | (32.0-50.0) | 12 | 38.5±6.3 | (28.0-48.5) | 17 |
| **QJT (pts)** | 32.3±4.5* | (23.0-41.0) | 14 | 30.5±3.4 | (24.0-38.5) | 11 | 27.6±3.8 | (20.5-33.5) | 14 |
| **SJ (cm)** | 35.1±6.2 | (25.5-47.4) | 18 | 33.1±5.7 | (24.5-48.8) | 17 | 32.6±8.8 | (22.6-47.1) | 27 |
| **CMJ (cm)** | 37.8±5.7 | (30.0-49.3) | 15 | 36.3±5.4 | (28.0-50.4) | 15 | 35.9±10.2 | (23.0-51.6) | 29 |
| **EI (%)** | 8.4±8.5 | (-2.6-30.2) | 101 | 10.4±6.3 | (-3.1-23.7) | 60 | 10.0±9.9 | (-1.5-30.6) | 99 |

**Note:** $\bar{X}$ = mean; SD = standard deviation; CV = coefficient of variation; QJT = quadrant jump test; SJ = squat vertical jump performance; CMJ = countermovement vertical jump performance; EI = elasticity index. \* $p < 0.05$ to *GUNGA*

Table 3 shows the values corresponding to the somatotype classification. Although there were differences in the mesomorphic component when comparing the VIOLA and MEDIO subgroups with the GUNGA subgroup (F_[2,47]_ = 7.617; p = 0.001) and ectomorphic component when comparing the VIOLA with GUNGA subgroup (F _[2,47]_ = 3.899; p = 0.027), the “endo-mesomorph” classification was found in the three subgroups. Fig 1 illustrates the predominance of the mesomorphic component among the two other components of the somatotype in the sample.

**Table 3.** Somatotype and its classification of the elite Capoeira athletes in the three categories (n = 50).

|  | VIOLA (n = 15) |  |  | MEDIO (n = 25) |  |  | GUNGA (n = 10) |  |  |
| --- | --- | --- | --- | --- | --- | --- | --- | --- | --- |
| | $\bar{X} \pm SD$ | Range | CV (%) | $\bar{X} \pm SD$ | Range | CV (%) | $\bar{X} \pm SD$ | Range | CV (%) |
| **Endomorphy** | 2.8±0.7 | (1.5-4.0) | 26 | 3.3±1.0 | (1.8-6.4) | 31 | 3.5±1.5 | (1.4-6.0) | 44 |
| **Mesomorphy** | 6.0±1.2* | (3.9-7.9) | 21 | 6.5±1.2* | (4.6-9.0) | 19 | 7.8±0.6 | (7.0-9.0) | 8 |
| **Ectomorphy** | 1.3±0.8* | (0.2-3.2) | 60 | 1.0±0.6 | (0.2-2.3) | 64 | 0.6±0.4 | (0.1-1.7) | 71 |
| **Classification** | Endo-mesomorph |  |  | Endo-mesomorph |  |  | Endo-mesomorph |  |  |

**Note:** $\bar{X}$ = mean; SD = standard deviation; CV = coefficient of variation. \* $p < 0.05$ to GUNGA

**Fig 1.**
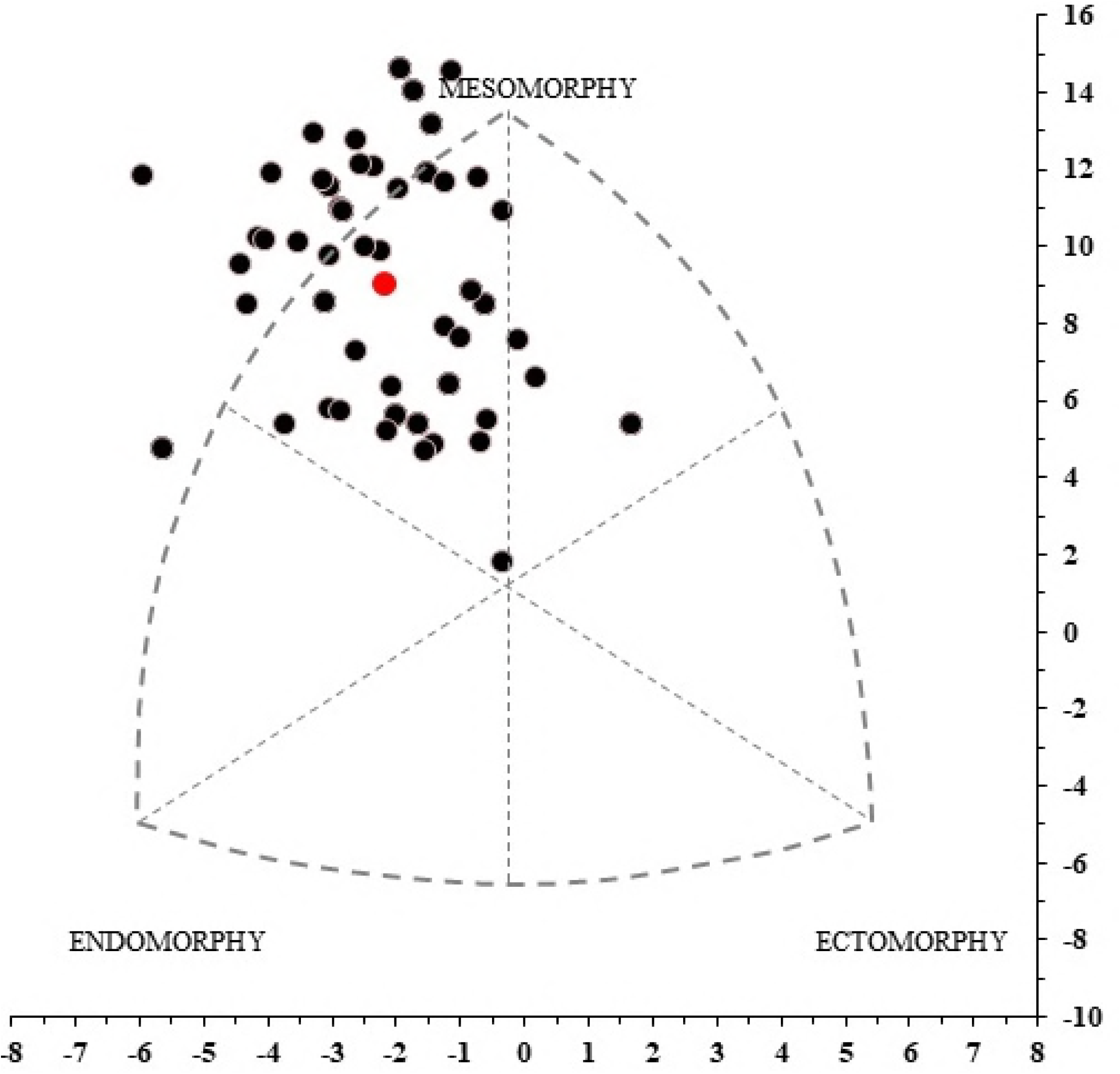
Somatochart of the elite Capoeira athletes. The intermediate red point represents the mean somatotype of the sample (n = 50).

Table 4 presents the correlation between somatotype and functional performance of elite Capoeira athletes. A negative correlation between the endomorphic component and performance in the sit-up (p < 0.001), push-up (p = 0.004), SJ (p = 0.001), and CMJ (p < 0.001) was found. The performance in the push-up correlated positively with mesomorphy (p = 0.009) and negatively with ectomorphy (p = 0.005).

**Table 4.** Relationship (r) between the anthropometric and functional variables of the elite Capoeira athletes in the three categories (n = 50).

|  | Endomorphy | Mesomorphy | Ectomorphy |
| --- | --- | --- | --- |
| **Push-up (reps)** | **-0.39\*** | **0.36\*** | **-0.39\*** |
| **Sit-up (reps)** | **-0.51\*** | -0.05 | 0.10 |
| **Sit-and-reach (cm)** | 0.14 | -0.05 | -0.01 |
| **QJT (pts)** | -0.17 | -0.16 | -0.07 |
| **SJ (cm)** | **-0.45\*** | 0.05 | -0.08 |
| **CMJ (cm)** | **-0.49\*** | 0.01 | -0.05 |

**Note:** QJT = quadrant jump test; SJ = squat vertical jump performance; CMJ = countermovement vertical jump performance. \*p < 0.05 to significant correlation

## Discussion

According to our knowledge, this is the first research to investigate anthropometric characteristics associated with functional performance of elite male competitors at the international level in Capoeira. From an anthropometric point of view, the main findings show that elite Capoeira athletes presented a predominance of the mesomorphic component, regardless of the weight subgroup in the competition (Table 3 and Fig. 1). Furthermore, the endomorphic component correlated inversely with the functional performance in the push-up, sit-up, CMJ, and SJ. However, the push-up showed a positive correlation with the mesomorphic component and a negative correlation with the ectomorphic component in the sample of elite Capoeira athletes (Table 4). These results reinforce the importance of a reduction in the relative BF added with the maintenance or gain of muscularity for someone that aims a great motor performance in a sport [26]. In addition, from a motor point of view, it was found that regardless of the weight subgroup in the competition, the functional performance did not present a difference between the VIOLA, MEDIO, and GUNGA subgroups (Table 2). All subjects in the sample had an excellent classification for the push-up, sit-up, and sit-and-reach test [23]. Although it was not the objective of this study, the results verified that the variables associated with general health and cardiovascular condition in the sample are within recommended levels, according to their BF [23], WHR [27], and systolic and diastolic blood pressure [28]. Moreover, controlled studies have demonstrated chronic benefits in the flexibility [8] and cardiovascular system [7] of Capoeira practitioners. These findings together support that the practice of Capoeira in medium (8–10 weeks) [7, 8] and long term (11–35 years of practice; Table 1) results into a favorable functional capacity profile and cardiovascular health of an individual.

In the combat sports perspective, the literature has documented the anthropometric profiles of jiujitsu athletes (n = 11) with a mean age of 25.8 ± 3.3 years [10], judo athletes (n = 104) aged 23.3 ± 3.0 years [11], taekwondo athletes (n = 146) [12], boxing fighters (n = 23) aged 19.3 ± 0.3 years [13], karate athletes (n = 19) aged 31.6 ± 8.8 years [14], and Greco-Roman wrestling athletes (n = 23) aged 24.9 ± 5.5 years [15]. Similar to these mentioned studies, the predominant somatotype classification in the present study was fitted in the mesomorphic component. The results found in the previously mentioned studies suggest that the long-term training program in combat sports operates in the modulation of the physical structure of the practitioner, which directs to the aspect of muscularity. As it was expected, absolute distinctions are demonstrated among these studies regarding relative BF. In elite Capoeira athletes (n = 50), this variable was observed in the mean of 11.4 ± 4.1%, while in karate, judo, jiujitsu, and taekwondo athletes, the values for BF were 14.7 ± 4.3%, 13.2 ± 2.5%, 10.3 ± 2.6%, and 9.6 ± 2.0%, respectively.

Referring to the endomorphic component of the somatotype, it was found that this variable showed an inverse relation with several functional tests in the sample of Capoeira athletes (Table 4). Abidin and Adam [29] investigated male and female martial arts athletes, and they demonstrated negative correlations between endomorphism and performance in tests, which assessed the explosive strength of lower limb muscles. These results demonstrate that fat accumulation in the central and peripheral regions may impair functional performance in capacities that require lower limb muscle strength. Fat accumulation also has a negative influence on the strength and resistance of the upper limb and abdominal muscles. Furthermore, combat sports commonly present mixed characteristics of metabolic requirement as in judo [30] and different rhythms of the game in Capoeira [31]. Ackland et al. (2009) [9] emphasize that fight sports require different physical qualities such as strength, endurance, speed, agility, and power, which are related to the mesomorphic component. This corroborates with the correlation found between mesomorphism and push-up (Table 4).

The height obtained in the performance of the CMJ in elite Capoeira athletes (36.7 ± 6.6 cm, Table 2) was inferior to those in taekwondo (40.8 ± 4.9 cm), jiu-jitsu (48.4 ± 4.8 cm), and judo (48.4 ± 2.9 cm) national athletes assessed in a noncompetitive period [32, 33]. In contrast, before an international jiujitsu competition, the athletes obtained a lower jump height (CMJ, 34.0 ± 5.2 cm), which is close to the result found in the present study [34]. Although specific movements in Capoeira and jiujitsu also require explosive qualities, such results may be associated with the training model of these martial arts. Moreover, the jumping test used may not be specific and sensitive enough to capture the best performance of these athletes. Future research on the frequency of use of these movements during noncompetitive games could contribute to the explanation of these results. The findings in this study are in line with those of other sports modalities, which affirmed that high-level athletes tend not to present a wide difference between the performance in CMJ and SJ [35, 36]. The low variation between the performance of these tests is supported by the EI of the different Capoeira athlete subgroups, which suggests adequate intermuscular coordination or a decrease in the muscle slack in the lower limbs [35]. However, further researches are still needed on the interrelationship of neuromuscular variables and a possible influence on athletic performance.

Capoeira is a modality that demands attack and defense movements, often with the practitioner being in upside-down positions, which demand physical qualities such as muscular power, muscular endurance, flexibility, and agility from different muscle groups [1, 37]. Considering this, the functional and anthropometric profiles presented may have practical application in the formulation of guidelines on a specific training program for the Capoeira practitioner who aims for a great performance within the modality. The results that can be achieved in physical composition and performance level from training programs will enable Capoeira practitioners to be in high-level competitive championships.

One limitation of the present study was not verifying the association of the anthropometric and functional variables with the real performance of the athletes in the competition. Therefore, further studies are needed to determine these relationships.

In conclusion, elite male competitors at the international level of the ABADÁ Capoeira School show predominance of the mesomorphic component in the somatotype. Furthermore, the endomorphic component correlated inversely with the functional performance of the trunk, upper limb resistance, and lower limb explosive strength. Further researches are needed to characterize the anthropometric and functional aspects of female athletes in the Capoeira modality, which still need to be clarified.

## Acknowledgement

The authors thank FACEPE (Research support foundation of the Brazil, Pernambuco State) to fund scholarships process IBPG-1203-4.09/16. The authors also thank Marcos Paulo Alves dos Santos (Prof. Gibor), Devanildo de Amorim Souza, Daniel Moraes (Inst. Fhel) and Patricia Bastos Indaê for the support provided in the data collection of this research.

